# Peptide-binding groove contraction linked to the lack of T-cell response: Using complex structure and energy to identify neoantigens

**DOI:** 10.1101/295360

**Authors:** Yuan-Ping Pang, Laura R. Elsbernd, Matthew S. Block, Svetomir N. Markovic

**Author notes:** Corresponding author: Stabile 12-26, Mayo Clinic, 200 First Street SW, Rochester, MN 55905, USA.

## Abstract

Using personalized peptide vaccines (PPVs) to target tumor-specific non-self antigens (neoantigens) is a promising approach to cancer treatment. However, the development of PPVs is hindered by the challenge of identifying tumor-specific neoantigens, in part because current in silico methods for identifying such neoantigens have limited effectiveness. Here we report the results of molecular dynamics simulations of 12 oligopeptides bound with a human leukocyte antigen (HLA), revealing a previously unrecognized association between the inability of an oligopeptide to elicit a T-cell response and the contraction of the peptide-binding groove upon binding of the oligopeptide to the HLA. Our conformational analysis showed that this association was due to incompatibility at the interface between the contracted groove and its αβ–T-cell antigen receptor (TCR). This structural demonstration that having the capability to bind HLA does not guarantee immunogenicity prompted us to develop an atom-based method 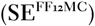 to predict immunogenicity through using the structure and energy of a peptide•HLA complex to assess the propensity of the complex for forming a ternary complex with its TCR. In predicting the immunogenicities of the 12 oligopeptides, 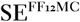 achieved a 100% success rate compared with success rates of 25–50% for 11 publicly available residue-based methods including NetMHC_-4.0._ While further validation and refinements of 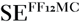 are required, our results suggest a need to develop in silico methods that assess peptide characteristics beyond their capability to form stable binary complexes with HLAs to help remove hurdles in using the patient tumor DNA information to develop PPVs for personalized cancer immunotherapy.

## I. INTRODUCTION

Immunotherapies that boost immunity against tumors are a promising approach to treating cancers [1,2]. Using personalized peptide vaccines (PPVs) [3] to activate T cells against tumor-specific non-self antigens caused by somatic non-synonymous mutations (neoantigens) is one such immunotherapy. Improved neoantigen identification combined with reverse immunosuppressive treatments such as PD-1 blockade could make this type of immunotherapy more specific, more effective, and safer than currently available immunotherapies [2,4,5].

The T-cell activation process involves the following steps [6-8]: A tumor-specific neoantigen binds to the peptide-binding groove of a class-I or class-II HLA after it is cleaved from a cancerous protein by the respective intra- or extracellular degradation. The neoantigen•HLA complex then translocates to the surface of an antigen-presenting cell and subsequently forms a ternary complex with its αβ–T-cell antigen receptor (TCR) that is a part of a multicomponent signaling system. The association of neoantigen•HLA with the TCR is then strengthened by the binding of co-receptor CD_4_ with class-II HLA or co-receptor CD8 with class-I HLA. The engaged TCR in turn signals another part of the multicomponent system, the CD_3_ complex. Ultimately, the CD_3_ complex triggers intracellular signaling cascades including the recruitment of kinases and the CD_4_/CD8-associated Lck for full activation. In this process, the binding of HLA with CD4 or CD8 is crucial both because it recruits the Lck and because the affinity of HLA for TCR is low (*K*_d_ = ~0.1–500 μM) relative to the nanomolar affinity complexation between antibody and antigen.

The identification of a neoantigen from patient tumor DNA is a formidable, multistep task that requires a search for an endogenous or synthetic peptide with the capabilities to form (*i*) a binary complex with an HLA, (*ii*) a ternary complex with the HLA and its TCR, and (*iii*) a quaternary complex with the HLA, TCR, and CD4 (for class-II HLA) or CD8 (for class-I HLA).

Two main categories of in silico methods have been developed to identify HLA-binding peptides as the first step of the multistep task. One category comprises residue-based (or sequence-based) methods such as those reported in Refs. [9-18]. These methods predict the HLA-complexation affinity or stability of a peptide with coarse granularity at the residue level. They require experimental HLA-binding data for “training” and use either the approximation that each residue of a peptide independently confers the affinity/stability of the peptide•HLA complex or the assumption that HLA-binding sequence pattern recognition assisted by machine learning algorithms can account for correlations among residues if the training dataset is reasonably large. Residue-based methods are computationally efficient and can perform a fast screen of a large number of peptide sequences, but they are ineffective in identifying peptides that bind class-II HLAs because these peptides have variable lengths and degenerate anchors [19].

The other category comprises atom-based (or structure-based) methods such as those reported in Refs. [20-29]. These methods are also known as “ab initio” methods because they do not require the experimental HLA-binding data. Atom-based methods predict the HLA binding affinity with fine granularity at the atom level and can be used to identify peptides that bind class-I and class-II HLAs. Atom-based methods assume that the native-state conformation of a peptide•HLA complex can be obtained by using a threading technique (*viz*., alignment of a query sequence of residues to a core template structure), a molecular docking technique, a physics-based molecular dynamics (MD) simulation technique, or combinations of these techniques if the objective function or forcefield used by these techniques is well parameterized and the configurational sampling is adequate. Herein, the objective function is a table of 20×20 predetermined statistical pairwise contact potentials that describes the compatibility between of an aligned query sequence of residues and its structural environment in a core template structure [30]; the forcefield is an empirical potential energy function with a set of predetermined parameters that describes the relationship between a structure and its energy [31]; the configurational sampling is either the sampling of various core template structures for their “fitness” to a query sequence of residues or the sampling of various conformations of the peptide•HLA complex. Both the threading and docking techniques are computationally inexpensive and can screen a large number of peptide sequences. The threading technique is ineffective in identifying peptides that bind HLAs with hydrophilic pockets because these hydrophilic structural environments differ from those of globular proteins upon which the objective function is based. The docking technique does not adequately account for the main-chain conformational flexibility of the peptide-binding groove in HLA and the effect of solvation or entropy on peptide binding at the groove. The MD simulation technique accounts for both the main-chain conformational flexibility and the solvation and entropy effects; however, it has been only rarely used for immunogenicity prediction because of its high computational cost that often limits the sample size to one MD simulation [32].

Owing to the approximations, assumptions, or known limitations of both categories of the current in silico methods, it is unsurprising that these methods have had limited success for neoantigen identification [33]. Further, we suspected that the limited success might be related to the uncertainty about whether HLA-complexation affinity/stability has a high, low, or no correlation with immunogenicity, even though immunogenic peptides are known to have the capability to bind HLAs. Some in silico methods were developed with an explicit assumption that the affinity or stability of a peptide•HLA complex is a correlate of immunogenicity of the peptide. Others were developed without any assumptions about this correlation. This ambiguity can lead to inappropriate attempts to identify immunogenic peptides using methods intended for the identification of HLA-binding peptides. If HLA binding affinity had a low or no correlation with immunogenicity, a new strategy would be needed to develop neoantigen identification methods that assess peptide characteristics beyond their capability to form stable binary complexes with HLAs.

Here we report our isobaric–isothermal MD simulations of 12 oligopeptides with known differences in eliciting a T-cell response (Table 1) [34,35]. These statistically rigorous simulations with an aggregate simulation time of 75.84 µs revealed a previously unrecognized association between oligopeptide-induced groove contraction in HLA and the inability of oligopeptides to elicit a T-cell response. We then report our conformational analysis of these simulations, showing that the association is due to incompatibility at the interface between the contracted groove of the binary complex and its TCR. This finding that HLA binding affinity has a low correlation with immunogenicity inspired us to develop an atom-based method 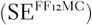 for neoantigen identification. 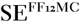 is conceptually new in that it uses the structure and energy of a peptide•HLA complex to assess the propensity of the binary complex for forming a ternary complex with its TCR. In this context, we also report the 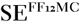 method and a comparison of 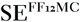 with 11 publicly available residue-based methods, including the NetMHC program, for retrospectively predicting T-cell responses to the 12 oligopeptides. Finally, we discuss a new strategy for the development of effective in silico methods to identify neoantigens as PPVs for personalized cancer immunotherapy.

**Table 1.**
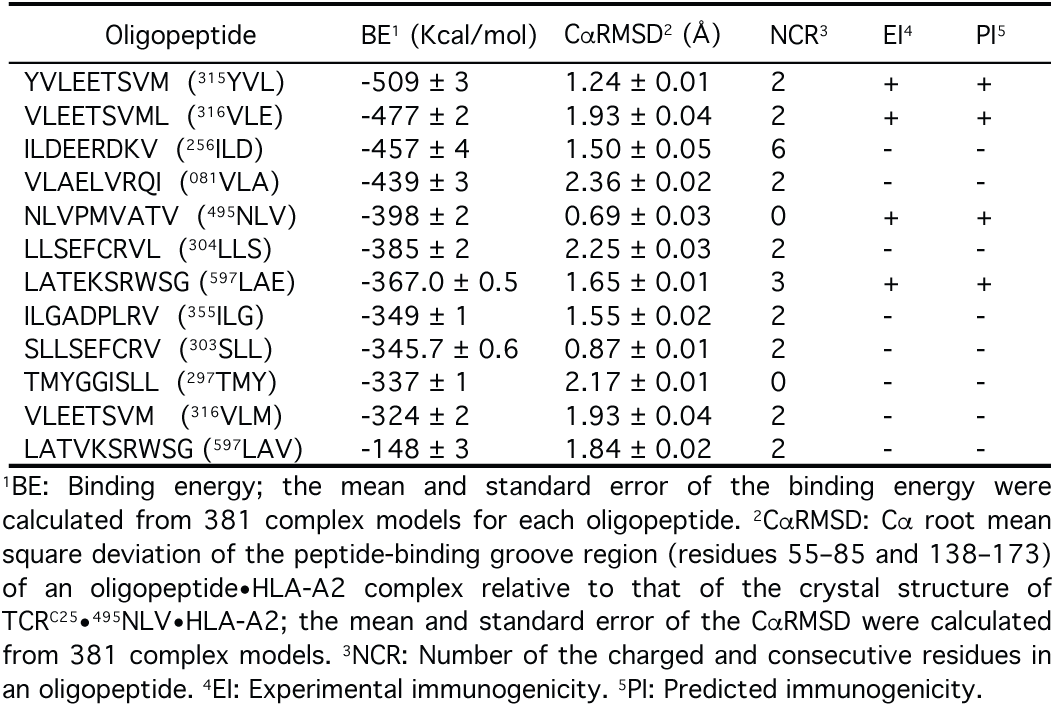
Predictions of immunogenicities of 12 oligopeptides using 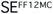.

## II. METHODS

### Model preparation

The initial conformation of a truncated class-I HLA A*02:01 (HLA-A2) liganded with an oligopeptide (residues 1– 183 of HLA-A2 and all residues of the oligopeptide) was taken from the crystal structure of HLA-A2 liganded with GVWIRTPPA (PDB ID: 5E00). GVWIRTPPA was mutated to each of the 12 oligopeptides listed in Table 1 using MacPyMOL Version 1.5.0 (Schrödinger LLC, Portland, OR). All D, E, R, K, and H residues in the 12 resulting complexes were treated as the AMBER residues of ASP, GLU, ARG, LYS, and HIE, respectively. All C residues in HLA-A2 and its ligand were treated as the AMBER residues of CYX and CYS, respectively. Each complex was then energy minimized using the SANDER module of AMBER 16 (University of California, San Francisco) with a dielectric constant of 1.0, a cutoff of 30.0 Å for nonbonded interactions, and 200 cycles of steepest descent minimization followed by 100 cycles of conjugate gradient minimization.

### MD simulations

Each energy-minimized complex was solvated with the TIP3P water [36] with surrounding counter ions for neutrality and then energy minimized for 100 cycles of steepest-descent minimization followed by 900 cycles of conjugate-gradient minimization using the SANDER module. The resulting system was heated from 5 K to 340 K at a rate of 10 K/ps under constant temperature and constant volume, then equilibrated for 10^6^ timesteps under constant temperature of 340 K and constant pressure of 1 atm employing isotropic molecule-based scaling, and finally simulated in 20 distinct, independent, unrestricted, unbiased, and isobaric–isothermal MD simulations with FF_12_MC [31] using the PMEMD module of AMBER 16 with a periodic boundary condition at 340 K and 1 atm. The numbers of TIP3P waters and surrounding ions and the initial solvation box size for each oligopeptide are provided in Table S1. The 20 unique seed numbers for initial velocities of Simulations 1–20 were taken from Ref. [37]. All simulations used (*i*) a dielectric constant of 1.0, (*ii*) Langevin thermostat [38] with a collision frequency of 2 ps^-1^, (*iii*) the Particle Mesh Ewald method to calculate electrostatic interactions of two atoms at a separation of >8 Å [39], (*iv*) Δ*t* = 1.00 fs of the standard-mass time [31,40], (*v*) the SHAKE-bond-length constraints applied to all bonds involving hydrogen, (*vi*) a protocol to save the image closest to the middle of the “primary box” to the restart and trajectory files, (*vii*) a formatted restart file, (*viii*) the revised alkali and halide ions parameters [41], (*ix*) a cutoff of 8.0 Å for nonbonded interactions, (*x*) the atomic masses of the entire simulation system (both solute and solvent) that were reduced uniformly by tenfold [31,40], and (*xi*) default values of all other inputs of the PMEMD module. The FF_12_MC parameters are available in the Supporting Information of Ref. [40]. All simulations were performed on a cluster of 100 12-core Apple Mac Pros with Intel Westmere (2.40/2.93 GHz).

### Conformational cluster analysis

The conformational cluster analysis of an oligopeptide•HLA-A2 complex was performed using CPPTRAJ of AmberTools 16 (University of California, San Francisco) with the average-linkage algorithm [42], epsilon of 2.0 Å, and root mean square coordinate deviation on all C*α* atoms of residues 55–85 and 138–173 of HLA-A2 and all Cα atoms of the ligand (see Table S2 for additional information). Centering the coordinates of the complex and imaging the coordinates to the primary unit cell were performed prior to the cluster analysis.

### Root mean square deviation and binding energy calculations

The Cα root mean square deviation (CαRMSD) for the truncated complex (residues 55–85 and 138–173 of HLA-A2 and all residues of the oligopeptide) or the truncated HLA-A2 (residues 55–85 and 138–173) was calculated using the ProFit program Version 2.6 (http://www.bioinf.org.uk/software/profit/). The binding energy was computed using the SANDER module and FF_12_MC with a dielectric constant of 1.0 and a cutoff of 30.0 Å for nonbonded interactions. For each oligopeptide, the mean and standard error of CαRMSD or binding energy were calculated from 381 complex models. These models include one model derived from 20 distinct simulations and 380 models derived from all combinations of 19 distinct simulations with one duplicate simulation. Namely, of the 381 models, the first was derived from {Simulations 1, 2, 3, … 19, 20}, the second from {Simulations 1, 1, 3, … 19, 20}, the third from {Simulations 1, 2, 1, … 19, 20}, …, the 21^st^ from {Simulations 2, 2, 3, … 19, 20}, …, and the 381^th^ from {Simulations 1, 2, 3, … 20, 20}.

### Success rate and robust Z score calculations

To avoid bias in calculating the success rate (SR) that is defined as the number of successful predictions divided by the number of peptides, the threshold for identifying active peptides was adjusted to obtain the maximal SR for each method (see Table S3 for adjusted thresholds). For example, the SR of NetMHC-4.0 was 30% when using the recommended percent rank threshold of 2%, and the maximal SR of 50% was obtained after changing the threshold to 3%.

The robust Z score of the *i*^th^ method [43] was calculated as below:

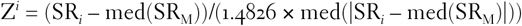

wherein median is the middle SR for an odd number of SR_s_ or the average of two middle SR_s_ for an even number of SR_s_; med(SR_M_) is the median of {SR_1_, SR_2_, … SR _*i*_, …, SR_M_}, *i* ∈{1, 2, …, M}, M is the number of distinct SR_s_ of N methods; med(|SR_*i*_ – med(SR_M_)|) is the median of {|SR_1_ – med(SR_M_)|, |SR_2_ – med(SR_M_)|, …, |SR_*i*_ – med(SR_M_)|, …, |SR_M_ – med(SR_M_)|}.

## III. RESULTS AND DISCUSSION

### Computer models of 12 oligopeptide•HLA complexes

To obtain the structural information that underlies a possible correlation between HLA binding affinity and immunogenicity, we computationally determined the most populated conformations (abbreviated as models) of HLA-A_2_ bound with the oligopeptides listed in Table 1. The decapeptides were obtained from the human B-Raf protein and its V600E neoantigen mutant [35]. All other oligopeptides were derived from the IE-1 and pp65 proteins of human cytomegalovirus [34]. These oligopeptides were chosen because of both their known differences in eliciting a T-cell response [34,35] and the reported crystal structures of ^495^NLV in complex with HLA-A2 alone or with HLA-A2 and TCR [44-46].

Because of the dependence of atom-based methods on forcefield and configurational sampling as noted in Section 1, we used our revised forcefield [31] in this study and applied our MD simulation method that uses uniformly reduced atomic masses to compress MD simulation time for improved configurational sampling [40]. FF_12_MC, which is a combination of our forcefield and low-mass simulation technique, has shown improved effectiveness in (*i*) autonomously folding fast-folding proteins, (*ii*) simulating genuine localized disorders of folded globular proteins, and (*iii*) refining comparative models of monomeric globular proteins [31]. Using FF_12_MC, we recently demonstrated that fast-folding proteins could be folded autonomously in isobaric–isothermal MD simulations to achieve agreements between simulated and experimental folding times within factors of 0.69–1.75 [47]. This outcome suggests that our forcefield and sampling technique are advanced and suitable for *a priori* determination of the most populated HLA-binding peptide conformation.

To address the high computational cost issue of MD simulations as noted in Section 1, we developed a customized, single-user cluster of 100 12-core Apple Mac Pros with which we can carry out 200 distinct, independent, isobaric–isothermal, and microsecond MD simulations simultaneously. In general, results derived from fewer than 20 MD simulations are considered unreliable [48,49]. In this study, we determined each HLA binary model from 20 distinct, independent, unrestricted, unbiased, isobaric–isothermal, 316-ns MD simulations.

Our previous studies showed that the most populated protein conformation of 20–40 distinct, independent, unrestricted, unbiased, isobaric–isothermal, sub-microsecond MD simulations resembled the experimentally determined native-state conformation for all of the proteins we had simulated using FF12-MC [31,47]. To confirm that our HLA complex models resemble the native-state conformations, we compared the ^495^NLV•HLA-A_2_ model that was derived *a priori* with corresponding crystal structures of ^495^NLV•HLA-A_2_, 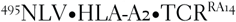, 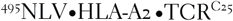, and 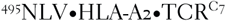. Of the 12 oligopeptides, ^495^NLV is the only one for which experimental complex structures have been reported (PDB IDs: 3GSO [44], 2×4R [45], 3GSN [44], 5D2N [46], and 5D2L [46]).

As indicated by CαRMSDs of 1.62 Å for 3GSO, 1.64 Å for 2×4R, 1.68 Å for 3GSN, 1.56 Å for 5D2N, and 1.62 Å for 5D2L, the peptide-binding groove region of the ^495^NLV model (residues 55–85 and 138–173 of HLA-A2 and all residues of ^495^NLV) resembled the corresponding region in all five crystal structures (Fig. 1). This finding supported our assumption that the native-state conformations of oligopeptide•HLA-A2 complexes could be obtained from our MD simulations using FF_12_MC, and hence we could investigate the correlation between HLA binding affinity and immunogenicity using the 12 HLA-A2 complex models, most of which lack reported experimental structures.

**Fig. 1.**
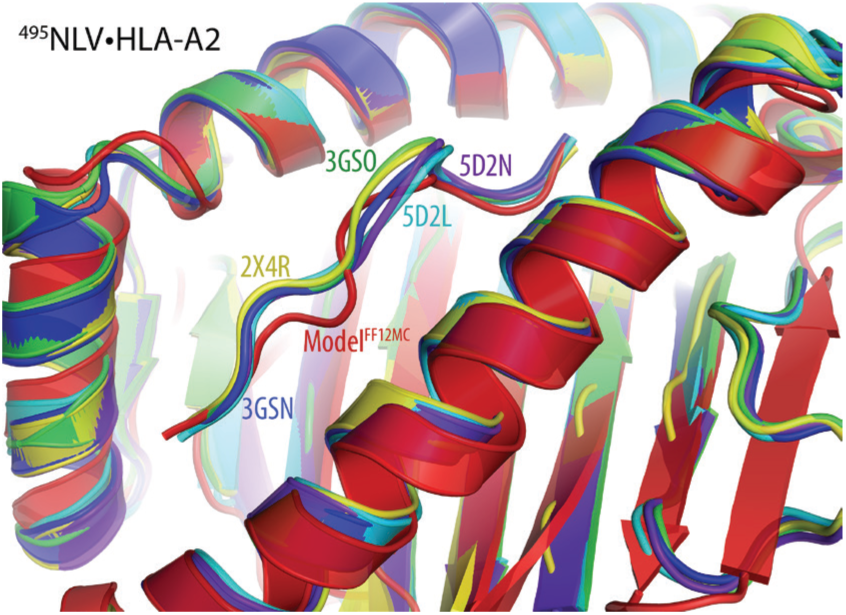
Overlay of the ^495^NLV•HLA-A_2_ model with corresponding crystal structures. The computer model was derived from 20 316-ns MD simulations using FF_12_MC. The five crystal structures are ^495^NLV•HLA-A_2_ (PDB IDs: 3GSO and 2×4R), 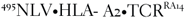 (PDB ID: 3GSN), 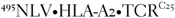 (PDB ID: 5D2N), and 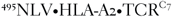 (PDB ID: 5D2L).

### Low correlation between affinity and immunogenicity

To investigate the correlation between HLA binding affinity and immunogenicity, we calculated the means and standard errors of the intermolecular interaction energies (abbreviated as binding energies) of the 12 complex models (Table 1). This type of binding energy indirectly accounts for the solvation and entropy effects that govern the most populated complex conformation because the binding energy was derived from the complex model—namely, from the most populated complex conformation of the 20 316-ns MD simulations using an explicit solvation model. It is therefore a reasonable proxy for binding free energy or binding affinity.

Of the 12 oligopeptides, only ^315^YVL, ^495^NLV, ^316^VLE, and ^597^LAE elicited a T-cell response [34,35]. If HLA binding affinity had a high correlation with immunogenicity, the binding energies of these oligopeptides would be significantly lower than those of the remaining eight oligopeptides. According to Table 1, only five oligopeptides had binding energies higher than those of the immunogenic ones. From the energy perspective, this suggests that HLA binding affinity has a low correlation with immunogenicity.

### Peptide-induced groove contraction in HLA

To further investigate this low correlation from the structure perspective, we analyzed the conformations of the 12 models from which the binding energies were derived. Notably, we observed that the models with ^256^ILD, ^303^SLL, ^315^YVL, ^316^VLE, ^355^ILG, ^495^NLV, and ^597^LAE adopted a common main-chain conformation (Fig. 2A) in the peptide-binding groove region (residues 55–85 and 138–173). This conformation was identical to the corresponding conformation of the 495NLV•HLA-A_2_ model and most resembled that of the crystal structure of 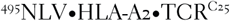 (PDB ID: 5D2N [46]). We also observed that the models with ^495^NLV, ^303^SLL, and ^315^YVL adopted a common main-chain conformation in the peptide segment (Fig. 2B), but the peptide sequences differed substantially. By contrast, the models with ^597^LAV and ^597^LAE had two markedly different main-chain conformations in the peptide segment (Fig. 2C) but the peptides differ by only one residue. Similarly, in their complex models, ^316^VLE and ^316^VLM had two drastically different backbone conformations (Fig. 2D) even though ^316^VLM is derived from 316VLE via a one-residue truncation at the *C*-terminus.

**Fig. 2.**
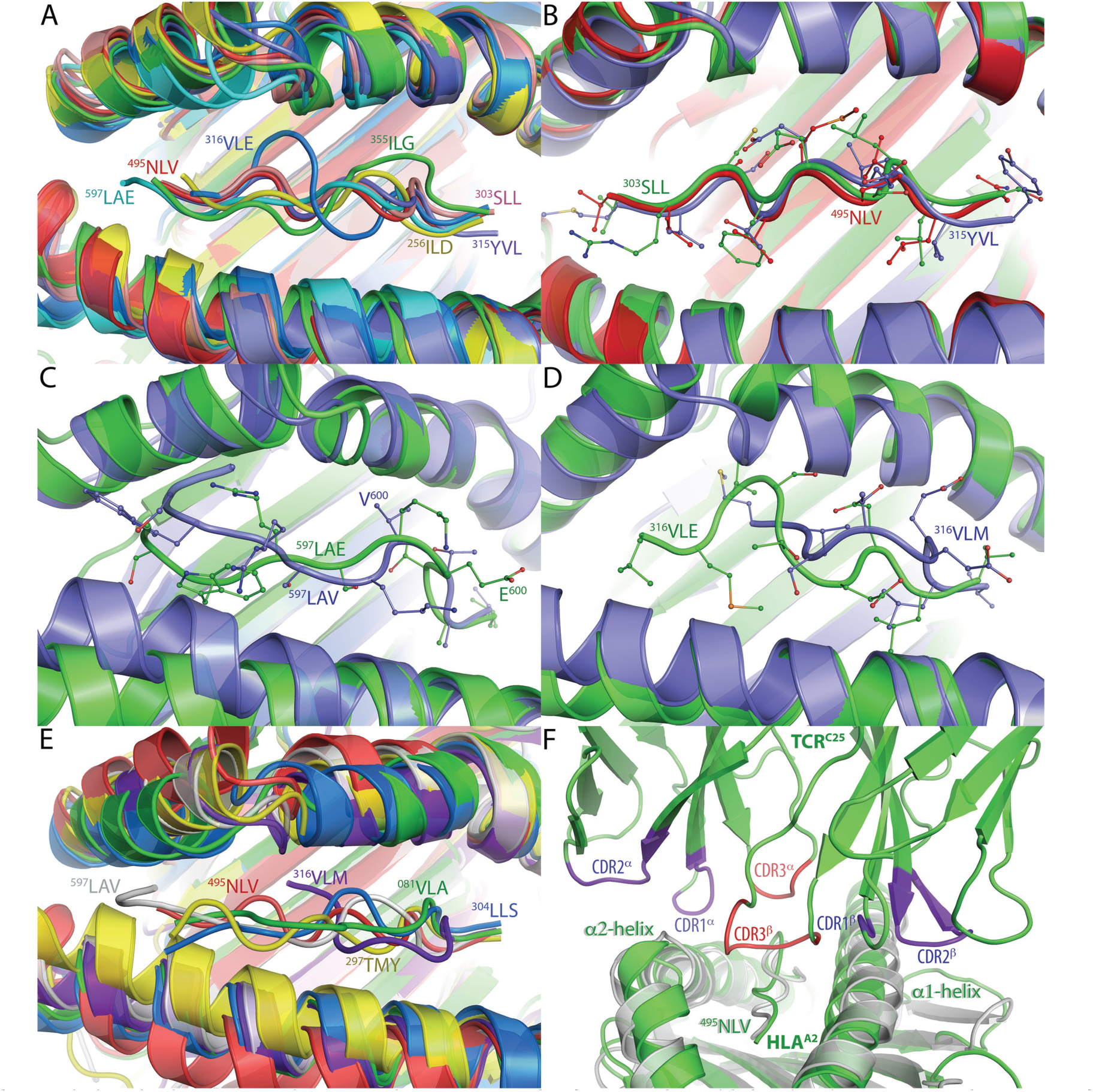
Oligopeptides bound with HLA-A_2_ or with HLA-A_2_ and C_25_ TCR. A. Overlay of seven complex models that adopted a common main-chain groove conformation. B. Overlay of three complex models that adopted a common main-chain oligopeptide conformation. C. Overlay of two complex models showing different main-chain oligopeptide conformations and the contracted groove conformation of ^597^LAV relative to that of ^597^LAE. D. Overlay of two complex models showing different main-chain oligopeptide conformations and the contracted groove conformation of ^316^VLM relative to that of ^316^VLE. E Overlay of six complex models showing five contracted groove conformations relative to that of ^495^NLV. F. Overlay of the ^495^NLV•HLA-A2 model onto the crystal structure of 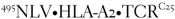 (PDB ID: 5D2N).

Unexpectedly, we found that the models with ^081^VLA, ^297^TMY, ^304^LLS, ^316^VLM, and ^597^LAV all had a contracted groove, with CαRMSDs of >2.1 Å for the groove region (residues 55–85 and 138–173) relative to that of ^495^NLV•HLA-A2. In other words, compared with that in ^495^NLV•HLA-A_2_ and other complexes, the space between the α1- and α2-helices in the five complexes was narrower (Fig. 2E). This contraction has an important implication for the correlation between HLA binding affinity and immunogenicity, as described below.

### Groove contraction linked to the lack of T-cell response

To investigate the groove contraction further, we evaluated the capability of the 12 complex models to form a ternary complex with a TCR and examined the relationship of this capability with the reported T-cell response. We found that all five peptides that induced the groove contraction (^081^VLA, ^297^TMY, ^304^LLS, ^316^VLM, and ^597^LAV; Fig. 2E) failed to elicit a T-cell response (Table 1). This association between the contraction of the peptide-binding groove induced by oligopeptide binding to HLA-A2 and the lack of T-cell response is consistent with current knowledge of the T-cell activation process, which involves multiple stepwise complexations of a peptide with HLA, TCR, and CD4/CD8 as described in Section 1.

When groove contraction occurs in a peptide•HLA complex, the formation of a ternary complex with its TCR requires groove expansion. At the cost of weakening the peptide interaction with the groove, this expansion is necessary to permit the interactions of conserved helical residues in the groove region with four canonical loops in the first two complementarity-determining regions (CDR1 and CRD2) of the α and β chains of a TCR [46] (Fig. 2F). These interactions dominate the TCR interface with the peptide•HLA complex and provide a conserved structural scaffold for the TCR to “screen” the sequences of peptides presented by an HLA [7,50]. The expansion is also necessary to accommodate the two loops in CDR3 of the TCR chains and enable their interactions with the central region of the peptide [46], which is required for the “pairing” of the TCR with its cognate antigen (Fig. 2F) [7,50]. However, because the peptide no longer binds tightly in the expanded groove, the screening of the peptide sequence and the TCR pairing are compromised if the groove is expanded. Therefore, a binary complex fails to form the ternary complex for eliciting a T-cell response when its groove is contracted to enable binding with HLA.

This previously unrecognized association demonstrates that having the capability to bind HLAs does not guarantee immunogenicity, although immunogenic peptides always have the capability to bind HLAs. This structural demonstration that HLA binding affinity is a component of immunogenicity may explain why HLA binding affinity has a low correlation with immunogenicity.

### Using complex structure and energy to identify neoantigens

The low correlation between HLA binding affinity and immunogenicity implied by our structural and energetic studies suggests a need to develop new methods for neoantigen identification that assess peptide characteristics beyond the capability to form a stable binary complex with a class-I HLA. Accordingly, we devised a new atom-based method 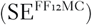 that uses the structure and energy of a peptide•HLA complex derived from MD simulations with FF_12_MC to assess the propensity of the complex for forming a ternary complex with a TCR. This method uses three predetermined energetic and structural thresholds.

As illustrated below, we were able to retrospectively predict the immunogenicities of the 12 oligopeptides using 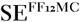 with (*i*) –360 kcal/mol as the binding energy threshold (to evaluate the capability of a peptide to form a binary complex), (*ii*) 2.00 Å as the CαRMSD threshold from the conformation of a ternary complex of HLA-A2 in the groove region (to assess the propensity of HLA-A_2_ in its binary complex to form a ternary complex), and (*iii*) four as a threshold for the number of charged and consecutive residues of an oligopeptide (to estimate the propensity of the peptide in its binary complex to form the ternary complex).

Of the ten cytomegalovirus oligopeptides, ^315^YVL, ^316^VLE, ^256^ILD, ^081^VLA, ^495^NLV, and ^304^LLS could be considered HLA-A_2_–binding peptides according to their binding energies (Table 1), which were lower than the threshold of –360 kcal/mol. However, the propensities of the ^081^VLA and ^304^LLS complexes for forming a ternary complex were lower than those of the rest of the six complexes according to their CαRMSD_s_ (Table 1), which were greater than the threshold of 2.00 Å. These larger CαRMSD_s_ imply that in the complexes of ^081^VLA and ^304^LLS, the narrow groove space has to expand to bind its TCR at the cost of weakening the peptide interaction with the groove. The propensity of the ^256^ILD complex for forming a ternary complex was also lower than those of all other complexes according to the number of charged and consecutive residues in ^256^ILD (Table 1), which was greater than the threshold of four. ^256^ILD has six charged and consecutive residues, and therefore the desolvation energy of its binary complex is too high to be compensated by the relatively weak interaction energies of the binary complex with its TCR. This conclusion is based on the fact that the *K*_d_ values of most TCR ternary complexes are in the low-affinity range of ~0.1–500 µM [7]. Therefore, only ^315^YVL, ^316^VLE, and ^495^NLV are likely to form both the binary and ternary complexes and be considered immunogenic. This retrospective prediction was in agreement with the experimental finding that only ^315^YVL, ^495^NLV, and ^316^VLE elicited a T-cell response [34].

Of the two B-Raf decapeptides, spontaneous CD8 T-cell responses to the neoantigen ^597^LAE were observed in HLA-A_2_^+^ patients with BRAF-mutant melanoma but not in patients with BRAF–wild-type melanoma or in healthy 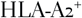 donors without melanoma [35]. Consistently, according to Table 1 and the three thresholds described above, even though the two peptides differ by one residue and ^597^LAV may bind HLA-A_2_ but elicit no T-cell response because it is a self-antigen, only ^597^LAE could be considered immunogenic.

### Comparison of 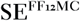 with residue-based methods

Compared with residue-based methods, 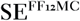 predicts HLA binding affinity at a finer granularity. Unlike residue-based methods that evaluate the propensity of a peptide to form a binary complex, 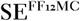 evaluates the propensities of a peptide to form both binary and ternary complexes. However, 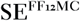 still does not analyze complexation with CD4 or CD8. Therefore, it is reasonable to question whether 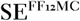 has a significant advantage over residue-based methods for identifying immunogenic peptides.

To evaluate the advantages of 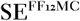, we retrospectively predicted T-cell responses to the 12 oligopeptides using 11 publicly available residue-based methods [9-18] and compared these predictions with those obtained with 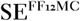 using the robust *Z* score standardization protocol to minimize the influence of outliers [43]. In this study the immunogenicity predictions with the 11 residue-based methods used a reported assumption that both the affinity and stability of an oligopeptide•HLA complex are a correlate of immunogenicity [17]. However, this assumption contradicts our results described above. Therefore, for the residue-based methods that are intended to identify HLA-binding peptides, their poor performances described below indicate misuses rather than deficiencies of the methods. Furthermore, we did not perform an exhaustive comparative study because the objective of the present work was to investigate the correlation between HLA binding affinity and immunogenicity for insight into new strategies to improve neoantigen identification and because we anticipated that refinements of 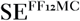 would be required to devise a method involving HLA, TCR, and CD4/CD8 that could ultimately be used in the clinic for personalized cancer immunotherapy.

Defining the success rate for each method as the number of successful predictions of T-cell response divided by the number of oligopeptides, we found that 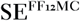 had a success rate of 100% and an outstanding robust *Z* score of 2.91, whereas all other methods had success rates of 25–50% and robust *Z* scores ranging from –1.13 to 0.22 (Table 2). These scores demonstrate that 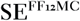 was significantly superior to the 11 residue-based methods for identifying the immunogenic oligopeptides in the present study. In addition, nine of the residue-based methods had the same success rate of 42% (Table 2), and eight of the nine methods including NetMHC-4.0 were reported as improvements to their predecessors. While further validation and refinements of 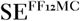 are required, the result of this comparative study suggests that combining the prediction of HLA binding affinity at a fine granularity with the prediction of capability of a peptide to form a ternary complex can significantly improve the identification of immunogenic peptides.

**Table 2.** Performance comparison of 12 in silico methods for prediction of immunogenicities of the 12 oligopeptides listed in Table 1.

| Method (reference) | Success rate (%) | Robust Z score |
| --- | --- | --- |
| $SE^{FF12MC}$ | 100 | 2.91 |
| NetMHCpan ([18]) | 50 | 0.22 |
| NetMHC-4.0 aka ANN ([16]) | 42 | -0.22 |
| BIMAS ([9]) | 42 | -0.22 |
| Consensus ([11,12,16]) | 42 | -0.22 |
| NetMHCcons ([15]) | 42 | -0.22 |
| NetMHCstabpan ([17]) | 42 | -0.22 |
| PickPocket ([14]) | 42 | -0.22 |
| SMM ([11]) | 42 | -0.22 |
| SMM<sup>PMBEC</sup> ([13]) | 42 | -0.22 |
| SYFPEITHI ([10]) | 42 | -0.22 |
| CombLib ([12]) | 25 | -1.13 |

### New strategy for developing neoantigen identification methods

In our comparison of 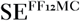 with the 11 publicly available residue-based methods for their capability to predict the likelihood of a T-cell response, 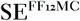 showed a significantly better accuracy (Table 2). More tellingly, the residue-based methods developed later showed no significant improvement over the BIMAS method that was reported in 1994 [9] (Table 2). These results underscore the inappropriate attempts to identify immunogenic peptides using methods intended for the identification of HLA-binding peptides and the problem of oversimplifying the multistep T-cell activation process into a single-step process. Further, the results suggest a need to expand the capability of 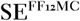 to predict immunogenicity with consideration of the involvement of HLA, TCR, and CD4/CD8.

Nevertheless, residue-based methods have long been preferred over atom-based methods because the latter are orders of magnitude more costly computationally. It is reasonable to question whether efforts should instead be directed toward expanding the capability of a residue-based method to account for peptide complexation with TCR and CD4/CD8. Given the limitations of current computing speeds, it is also fair to ask whether it is farfetched to aim for an atom-based method to predict immunogenicity with consideration of the involvement of HLA, TCR, and CD4/CD8.

As to the first question, a physical property descriptor of the number of charged and consecutive residues in a peptide for the partial evaluation of its likelihood to form a ternary complex can be easily implemented in residue-based methods. Because a cysteine-containing peptide can dimerize before binding to an HLA, another descriptor on whether a peptide contains a free cysteine can also be implemented such that ^304^LLS and ^303^SLL could be considered nonimmunogenic. These implementations would improve the performance of residue-based methods for immunogenicity prediction. However, it is difficult to incorporate the structural descriptor of the groove contraction in residue-based methods. Without this descriptor, as conceivable from the data in Table S3, residue-based methods would still be less effective than 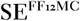 for assessing the propensity of a binary complex for forming a ternary complex, even if both the charge and cysteine descriptors were implemented, let alone the need to involve CD4/CD8.

As to the second question, it is worth noting that two strategies exist for developing computational methods to perform biomedical tasks. One strategy is to iteratively reduce the complexity of a codified task until the task can be carried out on an existing computer system. This type of codified task is often too simplified to be realistic, although this software-only strategy is relatively inexpensive because it does not require that engineers of computer hardware and software work together.

The other strategy is to iteratively develop a customized computer system with improved computing capacity until it can perform a codified task that is simplified yet reasonably realistic. This hardware-to-software strategy is costly because it requires computer hardware and software engineers to work together. Nevertheless, it is more suitable for a complicated task such as autonomous protein folding or neoantigen identification because the capacity of a customized computer system is confined only by the capital equipment budget for building the system. As an example, it was this hardware-to-software strategy that led to the development of computational methods capable of autonomously folding fast-folding proteins with explicit solvation [51]. Inspired by the pioneering work reported in Ref. [51], we developed a customized, third-generation, single-user cluster of 100 12-core Apple Mac Pros that enabled us to perform 240 isobaric–isothermal MD simulations with an aggregate simulation time of 75.84 μs for the present work.

It is also worth noting that in a few years instead of a decade well-controlled quantum computers are expected to be able to perform certain tasks faster than the most powerful transistor-based computers [52]. A 16-qubit quantum computer is now available for public use (https://quantumexperience.ng.bluemix.net/qx/devices) to promote the development of new in silico methods for performing exponentially scalable tasks relevant to neoantigen identification.

Given the success of the hardware-to-software strategy for the in silico folding of fast-folding proteins, the results of the present work, and the anticipation of quantum supremacy in five years or sooner, we suggest that the hardware-to-software strategy can be combined with some recent quantum computing techniques [53-55] to develop an advanced atom-based method for neoantigen identification. This method could reasonably account for cross-reactive epitopes [56] and use the energy and structure of a peptide•HLA•TCR complex to evaluate the propensity of the ternary complex for binding with CD4/CD8. Such a method may help overcome a considerable hurdle of using the patient tumor DNA information to develop PPVs for personalized cancer immunotherapy.

## Acknowledgments

This work was supported by the US Army Research Office (W911NF-16-1-0264 to YPP), the Mayo Foundation for Medical Education and Research, the Mayo Clinic Graduate School of Biomedical Sciences to LRE, and the Mayo Immunology Ph.D. Training Program to LRE. The content of this article is the sole responsibility of the authors and does not represent the official views of the funders.

### Author contributions

SNM, MSB, and LRE designed the overall immunogenicity prediction study using peptides derived from B-Raf and cytomegalovirus proteins; YPP conceived and implemented the 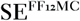 method and performed the immunogenicity prediction study using 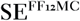. LRE performed the immunogenicity prediction study using 11 residue-based methods; YPP and LRE analyzed all prediction data; YPP wrote the paper; all authors contributed revisions.

### Competing financial interests

The authors declare no competing financial interests.

## Supplementary Information

Tables S1–S4.

